# A comparative analysis of computational tools for the prediction of epigenetic DNA methylation from long-read sequencing data

**DOI:** 10.1101/2021.04.24.441281

**Authors:** Shruta Sandesh Pai, Aimee Rachel Mathew, Roy Anindya

**Affiliations:** Department of Biotechnology, Indian Institute of Technology Hyderabad, Kandi, Sangareddy-502285, India

**Keywords:** DNA methylation, Long-read sequencing, Third-gen sequencing, Nanopore sequencing

## Abstract

Recent development of Oxford Nanopore long-read sequencing has opened new avenues of identifying epigenetic DNA methylation. Among the different epigenetic DNA methylations, N6-methyladenosine is the most prevalent DNA modification in prokaryotes and 5-methylcytosine is common in higher eukaryotes. Here we investigated if N6-methyladenosine and 5-methylcytosine modifications could be predicted from the nanopore sequencing data. Using publicly available genome sequencing data of *Saccharomyces cerevisiae*, we compared the open-access computational tools, including Tombo, mCaller, Nanopolish and DeepSignal for predicting 6mA and 5mC. Our results suggest that Tombo and mCaller can predict DNA N6-methyladenosine modifications at a specific location, whereas, Tombo dampened fraction, Nanopolish methylation likelihood and DeepSignal methylation probability have comparable efficiency for 5-methylcytosine prediction from Oxford Nanopore sequencing data.

## Introduction

Epigenetic DNA methylation primarily occur on cytosine or adenine. 5-methylcytosine (5mC) is the most ubiquitous DNA methylation present in eukaryotes ^[1]^, whereas, N6-methyladenine (6mA) is the most prevalent DNA modification in prokaryotes ^[2]^. However, high-resolution mass-spectrometry, 6mA was detected in some eukaryotes ^[3]^. Recently developed long-read PCR-free direct sequencing technology can also detect epigenetic DNA modification ^[4]^. Oxford Nanopore Technologies’ (ONT) nanopore sequencers quantifies the fluctuations of current when single-stranded nucleic acids pass through the nanopores and distinguishes the canonical bases from the 5mC or 6mA from the distinct patterns of variation of current ^[5]^. While a basecaller algorithm could detect the four canonical bases from the raw nanopore signals, the non-canonical bases can only be predicted using trained computational tools. In the present study, we compared computational tools that are available in public domains, namely, Tombo, mCaller, Nanopolish and DeepSignal for their efficiency to predict 6mA and 5mC modifications in *Saccharomyces cerevisiae* genomic DNA.

## Materials and Methods

### Sequence Data

We used ONT reads of Yeast MCM869 available on SRA database (run ID ERR2804505) that is part of the BioProject PRJEB28657. We downloaded the raw fast5 files using SRA-toolkit and is referred to as dataset 1 in further experiments. For dataset validation, we used the raw fast5 sequence data of yeast made available on DeepSignal GitHub repository by the authors of DeepSignal. This is referred to as dataset 2 in further experiments.

### Tombo

Tombo is a Mann-Whitney-U statistical comparison-based software tool, developed by ONT for the identification of non-canonical bases from nanopore sequencing data using Tombo command line interface. Tombo works on the principle of re-squiggling raw nanopore reads, where the squiggled raw nanopore signal is aligned to the reference genome. In our study, Tombo was used for the prediction of both 5mC and 6mA methylation in *S. cerevisiae* genome ^[6]^. The latest available version of Tombo (1.5.1) was installed on Linux OS via bioconda environment (on python 3.6 support). The re-squiggle algorithm takes an input read file (.fast5 file format) containing raw nanopore signal and associated base calls (with the help of a basecaller) along with a reference genome. Re-squiggling was the first step carried out by Tombo and the sequence to signal assignment was saved back into the read .fast5 file format. This step created a hidden file alongside the fast5s directory containing the essential genomic location for each read. The next step was to detect modifications using Mann-Whitney U statistical test and this generated a binary statistics file in .hdf5 file format and contained statistics associated with each genomic base producing a valid result. The following steps in the Tombo downstream pipeline made use of the above generated binary .stats file as an input. The ‘tombo text_output browser_files’ command generated output text files for both plus and minus strands in wiggle file format for the dampened fraction and bedGraph file format for the coverage, which can be used for visualization in the Integrated Genome Viewer (IGV) and Circos plot, and for calculation of ratio of methylated motifs to unmethylated motifs. Finally, the most significant modified base positions were obtained from the raw signal in the form of a plot by running the command ‘tombo plot most_significant’.

### mCaller

mCaller is a neural network-based python software tool that can predict DNA 6mA methylation from nanopore signal data and it works on the principle of predicting 6mA methylation based on the differences between measured and expected currents ^[7]^. The latest available version of mCaller (1.0) was installed on Linux OS directly from GitHub url and the dependencies/requirements for mCaller (bwa mem, scikit-learn, h5py, biopython, matplotlib, seaborn, numpy, pysam, scipy, pandas) were installed via conda. This program algorithm takes an input read file (.fast5 file format) containing raw nanopore signal and associated base calls and a reference genome and predicts methylations in GATC motifs based on a pre-existing model. Firstly, the template strand reads were extracted from the .fast5 files and then saved to a file path in .fastq file format using the ‘nanopolish extract’ command. These reads were then indexed using ‘nanopolish index’ command. The .fastq reads, which were obtained after the extraction step, were aligned with the reference genome using bwa mem to produce a sorted bam file. Prior to the prediction of DNA 6mA modification, the events values were scaled towards the model using the ‘nanopolish eventalign’ command to generate a .tsv file. The prediction of DNA 6mA modification step by mCaller returned a tabbed file with per-read predictions, where the columns indicated the chromosome number, genomic location, strand identity, label and methylation probability predicted by mCaller for that position and read. The last step in the methodology pipeline was to generate a summary methylation text .bed file of per-position methylation predictions. This output file in the .bed file format was converted to the bedGraph file format for proper visualization in IGV and Circos plot, and for calculation of ratio of methylated motifs to unmethylated motifs.

### Nanopolish

Nanopolish is based on Hidden Markov Model (HMM) which computes the probability of observing a modified base (5mC) based on the differences in the event distributions of methylated and unmethylated DNA ^[8]^. The latest available version of Nanopolish (0.9.0) was installed using ‘apt install’ along with the other prerequisites namely samtools and minimap2. This program algorithm takes an input read file (.fast5 file format) containing raw nanopore signal and associated base calls and a reference genome. We first ran the ‘extract’ command to extract basecalled information from fast5 files into an output .fastq file. This file was further indexed using ‘index’ command. With the help of minimap2 and samtools, the .fastq file was aligned to a reference .fasta file for the genome to produce a .bam file that was further indexed. Next, methylations were predicted with the bam file as input using the ‘call-methylation’ command. Output was obtained as .tsv file containing log_likelyhood_methylated, log_likelyhood_unmethylated and log_likelyhood_ratio for a particular position in every read. Lastly, a methylation summary file was obtained using the python script available to give methylation frequency and coverage per position. Using this output, bedGraph files for log_likelyhood_ratio, methylation frequency and coverage were created which were further analyzed with IGV and Circos plot and used for the calculation of ratio of methylated motifs to unmethylated motifs.

### DeepSignal

DeepSignal is a neural network-based tool that employs two modules to construct features from raw electrical signals of ONT reads ^[9]^. This is performed by using the convolutional neural network (CNN) to construct features directly from raw electrical signals followed by the bidirectional recurrent neural network (BRNN) to construct features from sequences of signal information which is then together fed into a fully connected neural network to predict the 5mC methylation states ^[9]^. The predictions using DeepSignal were performed using google colaboratory platform by installing DeepSignal (0.1.8) with pip and other prerequisites as well namely python (3.6.0) with conda, ont-tombo with conda and tensorflow (1.12) with pip. The trained model for 5mC in the context of CpG motif was downloaded from google drive link available on GitHub repository. We first ran the ont-tombo ‘resquiggle’ command on the fast5 data to re-squiggle the input based on a reference genome file followed by the deepsignal ‘extract’ command to extract the signal features of motifs of defined length (17-mer) and sequence (CG). These motif features were saved as .tsv file which was further used to call 5mC modifications using ‘call_mods’ command. This command produced .tsv file as output that included probability_0 (unmethylated), probability_1 (methylated) and called_label which was either 0 or 1 based on the probabilities for each position in every read. Lastly, a methylation summary file was obtained by running the python script available to give methylation frequency and coverage per position. Using these outputs, bedGraph files for probability_0, probability_1, methylation frequency and coverage were created which were further analyzed with IGV and Circos plot and used for the calculation of ratio of methylated motifs to unmethylated motifs. The details of all commands used to install and run the computational tools are provided in Supplementary Information.

## Results and Discussion

### Prediction of 6mA modification by Tombo and mCaller

Although 6mA is not common in eukaryotes, it was shown to be present in *S. cerevisiae* and suggested to have regulatory role in transcription elongation ^[10]^. When PacBio sequencing was used to sequence *S. cerevisiae* genome, several sites for 6mA were predicted ^[11]^, however, this data has been shown to be a result of overestimation due to presence of artifacts ^[12]^. We first examined Tombo tool for 6mA prediction. The successful completion of Tombo tool generated outputs in the form of three file formats-wiggle, bedGraph and pdf file formats. The output .wig text files, for both the forward and reverse strands, gave information regarding the base position and its dampened fraction i.e. the estimated fraction of modified bases in the context of GATC motif at that position. We converted Tombo output .wig files into .bedgraph file format and utilized the bedGraph files from Tombo and mCaller tools to create configuration files for generating Circos plot. The Circos plot gave *S. cerevisiae* genome view and chromosome XII had a significantly high methylation peak from Tombo as shown in **Figure 1A (ii and iii)** for forward and reverse strands respectively. The .wig files were then loaded as tracks into IGV (with SacCer3 as the genome to be compared with) for visualizing the methylation patterns and certain parameters such as track height, track color and windowing function were adjusted for better visualization purposes. Another parameter ‘data range’ was set to a minimum of 0.5 to eliminate the redundant methylation values. Therefore, after adjusting all the required parameters, we were clearly able to observe a high methylation peak on chromosome XII for both the strands of *S. cerevisiae*. We focused specifically on the chromosome XII using IGV and the significant high methylation peak was observed at the approximately 1 Mb region on the right arm of chromosome XII known as ribosomal DNA (*RDNA*) locus from Tombo (**Figure 1B, i and ii**) for forward and reverse strands. The output bedGraph files, for both the forward and reverse strands, gave information regarding the base position and its coverage i.e. the number of unique reads that include a given modified base in the context of GATC motif during the nanopore sequencing. These bedGraph files were then loaded as tracks into IGV and a high coverage density was observed at the *RDN* locus on chromosome XII of *S. cerevisiae* as shown in **Figure 2A (ii and iv)**. The output pdf plot was obtained by aligning the basecalled nanopore data to the reference genome as shown in **Figure 2B**. This plot gave information regarding the most significant modified base positions, strand, estimated fraction alternate and coverage.

**Figure 1.**
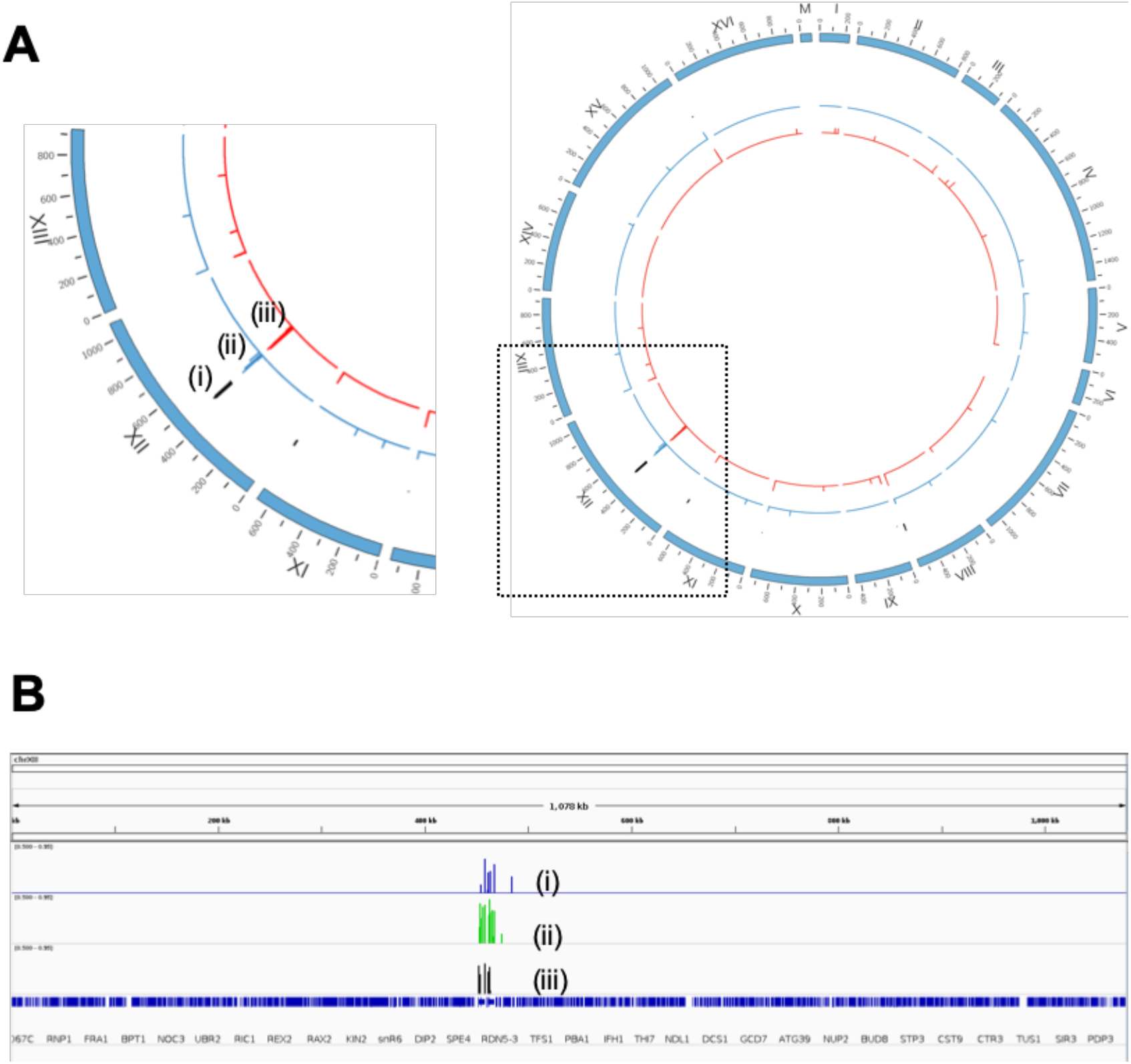
Prediction of DNA 6mA modification by mCaller and Tombo. **A**. Prediction of significantly high 6mA modification using Circos plot on *S. cerevisiae* genome showing all chromosomes in a circular map with ticks at every 100kbp (i) Histogram track showing DNA 6mA prediction by mCaller. Histogram track showing DNA 6mA prediction in the forward strand (II) and the reverse strand (iii) by Tombo. **B**. Analysis of the prediction of high 6mA methylation peak at the *RDN* locus on chromosome XII using IGV. On top, the region of chromosome XII with its size is written below it and on the bottom, the expanded gene view is displayed. (i) Track showing high DNA 6mA prediction by Tombo, for forward strand, at the locus (threshold >0.5). (ii) Track showing high DNA 6mA prediction by Tombo, for reverse strand, at the locus (threshold >0.5) (iii) Track showing high DNA 6mA prediction by mCaller at the locus (threshold >0.5). The box indicates region of high dampened fraction.

**Figure 2.**
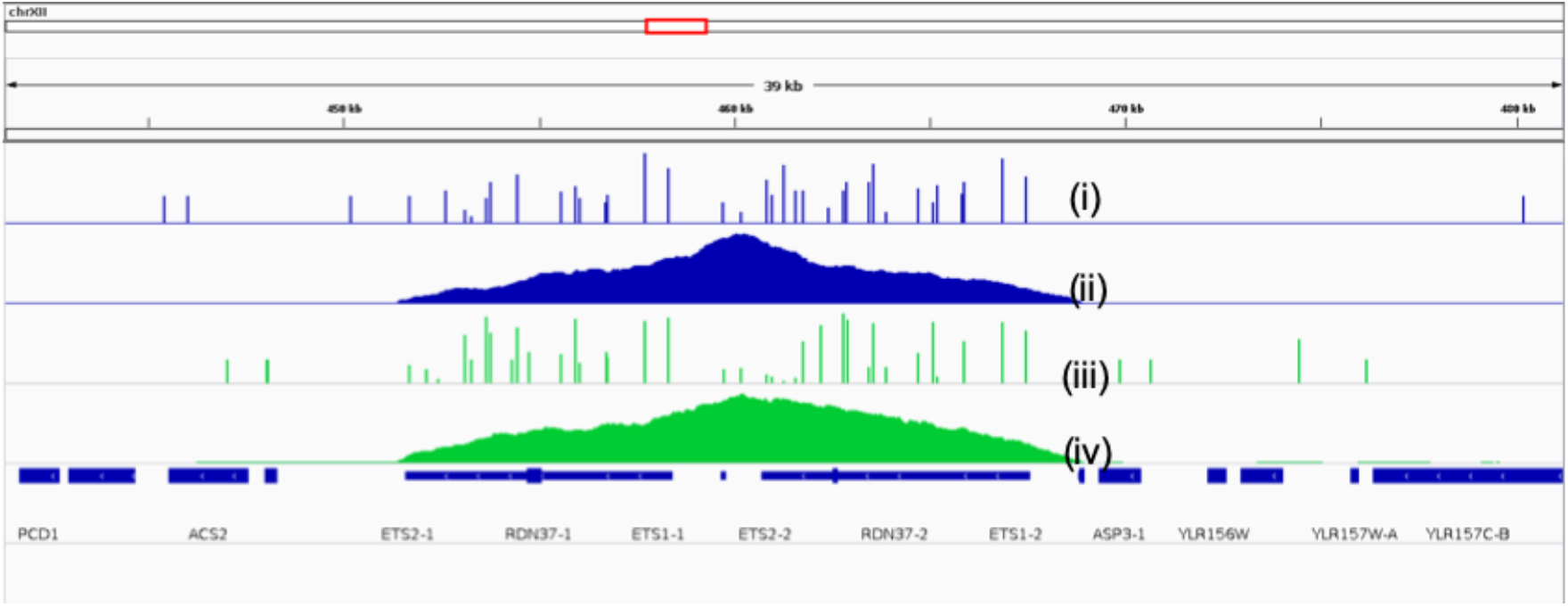
Prediction of high 6mA methylation at the *RDN1* locus by Tombo. The red rectangular box on top indicates the region of chromosome XII that is written below and on the bottom, the expanded gene view is displayed. (i) Track showing high 6mA prediction for forward strand (threshold >0.5). (ii) Track showing high coverage for 6mA prediction in forward strand. (iii) Track showing high 6mA prediction for reverse strand (threshold >0.5). (iv) Track showing high coverage for 6mA prediction in reverse strand.

Next, we examined mCaller tool for 6mA prediction. The goal was to rule out error regarding 6mA prediction using Tombo tool. The successful completion of mCaller tool generated methylation summary output in the form of .bed file format on applying the minimum read depth for inclusion as 7 and the minimum fraction of methylated base position as 0.5, i.e., when the predicted frequency of methylation was greater than 50%. The output .bed file gave the necessary per-position methylation predictions, and the .bed file was later converted to .bedgraph file format. The same required parameters were adjusted for the bedGraph track loaded into IGV and we were able to observe a significantly high methylation peak at the *RDN* locus on chromosome XII of *S. cerevisiae* as shown in **Figure 1B (iii).** For dataset validation, we used the raw fast5 data from nanopore sequencing of yeast made available on the DeepSignal GitHub repository containing ~4000 R9.4 1D reads basecalled by Albacore (dataset 2). We obtained outputs consistent with dataset 1 with a high coverage peak at *RDN* locus on chromosome XII. Within the locus, we observed significantly high 6mA methylation peaks from Tombo and mCaller and was consistent with dataset 1. The results for validation experiments are available in the supplementary information.

The output derived for 6mA from mCaller provided predictions for methylation of GATC motif for every position in every read. Based on a threshold set, these positions were called to be methylated or unmethylated and the frequency of methylation for called sites along with coverage was provided for each position whereas Tombo for 6mA, in the context of GATC motif, provided dampened fraction directly that gave methylation prediction per position after considering the repeated reads. For quantitative analysis **(Table 1),** we summarized the total number of motifs that were read to be present and from the prediction output, we derived how many of those motifs were predicted to be methylated according to each computational tool and its ratio was calculated. We performed this analysis for the whole genome as well as for chromosome XII. We observed that for the whole genome in the case of 6mA, the number of GATC motifs was much lower as compared to number of CpG motifs. We observed that mCaller read more GATC motifs and gave a higher percentage of methylation prediction of 3.14% when the threshold of probability was set to 0.8 compared to that of Tombo which read lesser GATC motifs and gave a lower percentage of methylation prediction of 0.88%. In case of chromosome XII, Tombo prediction percentage was 6.53% which is much higher than its whole genome equivalent with increase of over 8-fold whereas mCaller prediction percentage was almost the same as its whole genome equivalent. Despite their varying detection principles of non-canonical bases, both Tombo and mCaller produced outputs that supported the presence of 6mA modification in *S. cerevisiae* genome. As per the IGV analysis of 6mA dampened fraction file from Tombo and 6mA methylation summary file from mCaller, we obtained consistently significant methylation peaks at the *RDN* locus on chromosome XII. The *RDN* locus not only harbors the repeated units encoding ribosomal RNAs (rRNA), but also is the site of DNA replication, transcription and recombination ^[13]^. Here, approximately 150 repeated units are tandemly arranged and each single unit of contains two genes, the RNA pol I- transcribed 35S rRNA gene, and RNA pol III-transcribed 5S rRNA gene. The 35S rRNA is further processed to generate the 18S, 5.8S and 25S rRNAs. Therefore, the prediction of 6mA in the *RDN* locus is intriguing. Given the role of 6mA in pausing RNA pol II elongation ^[10]^, whether 6mA modification facilitates active transcription of RNA pol I and pol III remains to be explored.

**Table 1.**
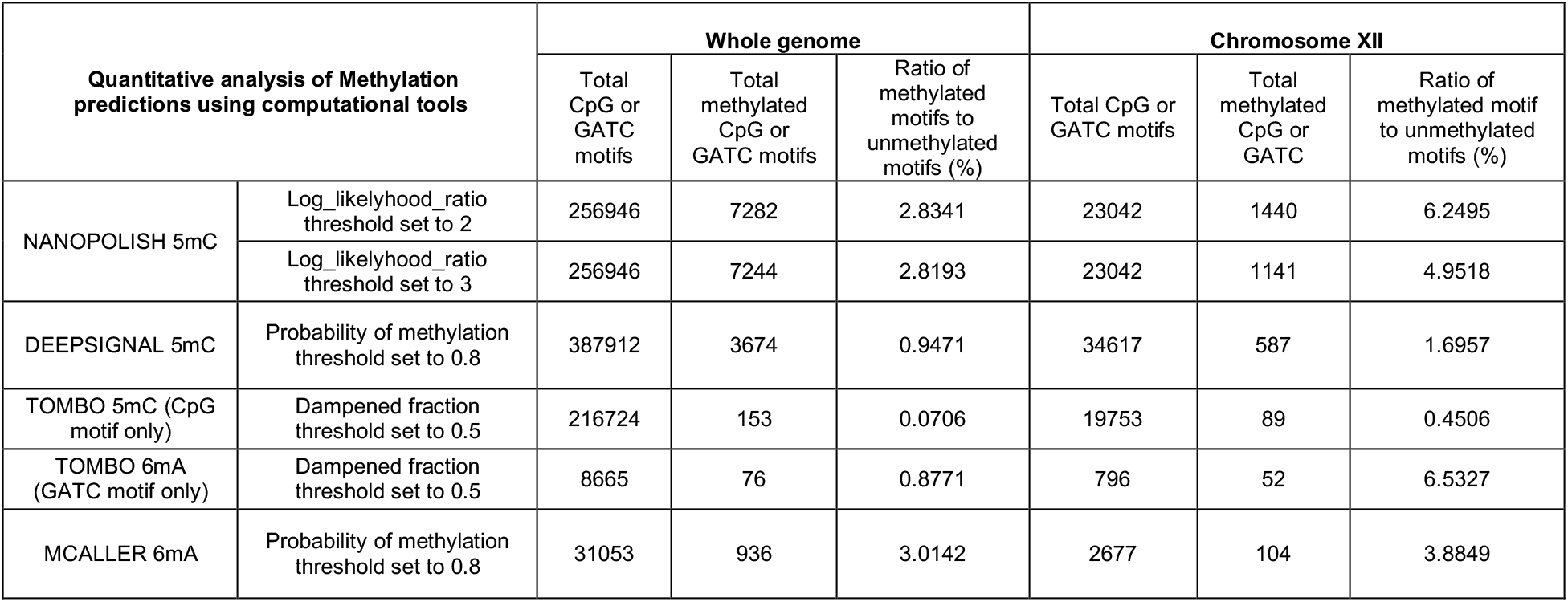
Summary of quantitative analysis of methylation using computational tools. The analysis was performed for the whole genome as well as for chromosome XII.

| Quantitative analysis of Methylation predictions using computational tools |  | Whole genome |  |  | Chromosome XII |  |  |
| --- | --- | --- | --- | --- | --- | --- | --- |
|  |  | Total CpG or GATC motifs | Total methylated CpG or GATC motifs | Ratio of methylated motifs to unmethylated motifs (%) | Total CpG or GATC motifs | Total methylated CpG or GATC | Ratio of methylated motif to unmethylated motifs (%) |
| NANOPOLISH 5mC | Log_likelihood_ratio threshold set to 2 | 256946 | 7282 | 2.8341 | 23042 | 1440 | 6.2495 |
|  | Log_likelihood_ratio threshold set to 3 | 256946 | 7244 | 2.8193 | 23042 | 1141 | 4.9518 |
| DEEPSIGNAL 5mC | Probability of methylation threshold set to 0.8 | 387912 | 3674 | 0.9471 | 34617 | 587 | 1.6957 |
| TOMBO 5mC (CpG motif only) | Dampened fraction threshold set to 0.5 | 216724 | 153 | 0.0706 | 19753 | 89 | 0.4506 |
| TOMBO 6mA (GATC motif only) | Dampened fraction threshold set to 0.5 | 8665 | 76 | 0.8771 | 796 | 52 | 6.5327 |
| MCALLER 6mA | Probability of methylation threshold set to 0.8 | 31053 | 936 | 3.0142 | 2677 | 104 | 3.8849 |

### Prediction of 5mC modification by Tombo, nanopolish and DeepSignal

Although the importance of 5mC as epigenetic DNA modification is well known in mammalian cells, it is considered to be absent in *S. cerevisiae*. However, using a sensitive method based on gas chromatography/mass spectrometry (GC/MS), it was recently shown to be present at very low concentration in several yeast species, including *S. cerevisiae* ^[14]^. Whole-genome sequencing of *S. cerevisiae* with PacBio sequencing revealed that yeast genome has about forty 5mCs and thousands of 6mAs [11]. Having predicted the enrichment of 6mA in *RDN* locus of yeast genome, we evaluated Tombo tool for 5mC prediction in the context of CpG motifs. On IGV analysis of .wig files with threshold of 0.5 for dampened fraction, we observed few low peaks in some of the telomeric regions of yeast chromosomes and a high peak in the middle of chromosome XII as shown in **Figure 3A (i** and **ii)** for forward and reverse strands respectively. The bedGraph files with coverage showed high values corresponding to the peak observed for dampened fraction as shown in **Figure 3A (iii** and **iv)** for forward and reverse strands respectively. On further analysis, it was observed that the peak represented *RDN* locus of the genome as shown in **Figure 4A (i** and **ii)**. Notably, the high coverage at the locus could probably be due to the repetitive nature of the locus. It was also observed that methylations were present on both forward as well as reverse strand of the *RDN* locus and in coding as well as non-coding region with no positional bias to regulatory elements.

**Figure 3.**
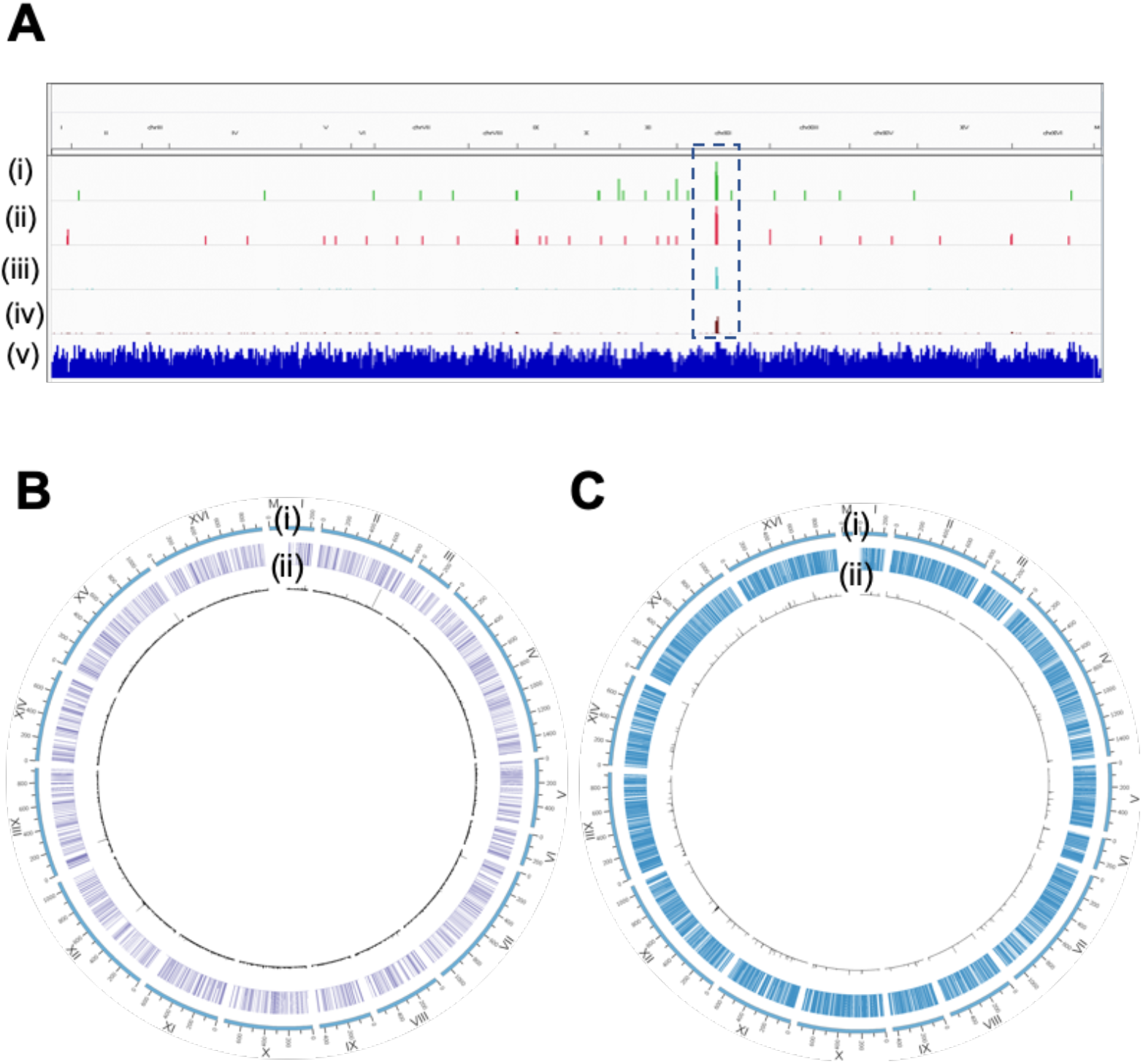
Prediction of 5mC methylation sites by Tombo, NanoPolish and DeepSignal. **A**. *S. cerevisiae* full genome sequence as viewed on IGV with tracks from Tombo output (i) Dampened fraction of 5mC on the positive strand (green) (threshold >0.5), (ii) Dampened fraction of 5mC on the negative strand (red) (threshold >0.5), (iii) Coverage on positive strand, (iv) Coverage on negative strand and (v) Genes. The box(- - -) indicates region of high dampened fraction as well as coverage **B**. Circos plot for Nanopolish 5mC prediction in *S. cerevisiae* genome showing all chromosomes in a circular map with ticks at every 100kbp and histogram tracks for each output data (i) Methylation Frequency at each base (threshold >0.5) (ii) Positive values of Log likelihood ratio of the base being methylated in a particular read. (0 to 11) **C.** Circos plot for DeepSignal 5mC prediction in *S. cerevisiae* genome showing all chromosomes in a circular map with ticks at every 100kbp and histogram tracks for each output data (i) Methylation Frequency at the base (threshold >0.5) (ii) Probability of the base being 5mC in a particular read (threshold >0.9)

**Figure 4.**
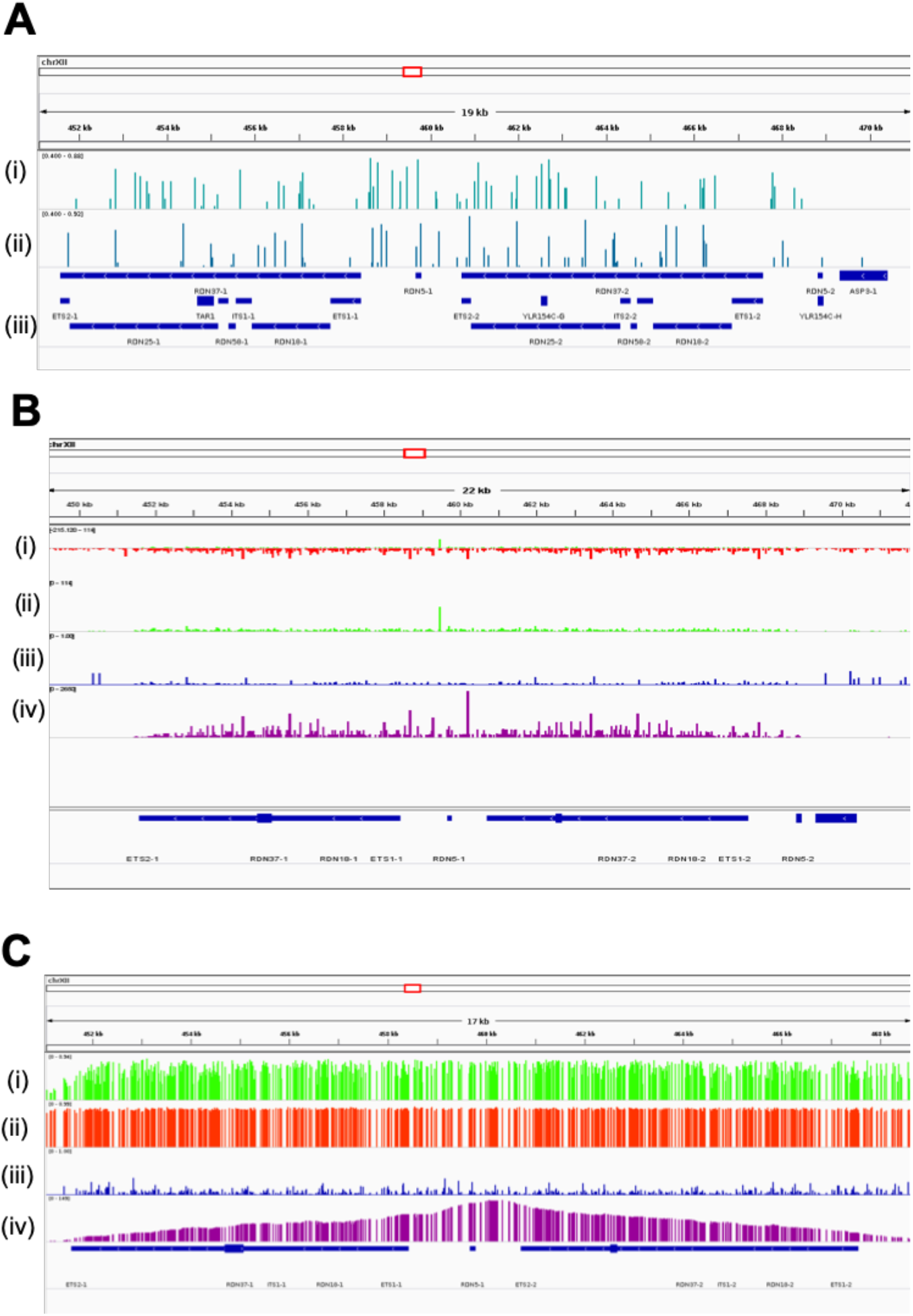
Prediction of 5mC methylation at the *RDN* locus on *S. cerevisiae* chromosome XII. For all figures, the red rectangular box on top indicates the region of chromosome XII that is zoomed in with its size written below. **A.** IGV analysis of Tombo output (i) Dampened fraction of 5mC methylated residues on the positive strand (0.4 to 0.88), (ii) Dampened fraction of 5mC methylated residues on the negative strand (0.4 to 0.92) and (iii) Genes. **B.** IGV analysis of Nanopolish output (i) Log likelihood ratio of 5mC methylation with positive values in green and negative values in red (−215 to 11) (ii) Log likelihood ratio of 5mC methylation with positive values only (0 to 11) (iii) Methylation frequency of 5mC (blue)(0 to 1) (iv) Read Coverage. **C.** IGV analysis of DeepSignal output (i) Probability of being 5mC methylated (0 to 0.94) (ii) Probability of being unmethylated C (0 to 0.99) (iii) Methylation Frequency of 5mC (0 to 1) and (iv) Coverage.

Nanopolish computational tool gave output in terms of log likelihood ratio of 5mC methylation in the context of CpG motifs and was used for predicting DNA methylation ^[11]^. Positive values of the ratio indicated positions that are likely to be methylated and negative values indicating otherwise. We observed that very few regions are predicted to have high positive values of the likelihood ratio for 5mC methylation, *RDN* locus being one such region as shown in **Figure 3B (ii)**. The summary python script to calculate methylation frequency considered all the reads having positive value of the ratio to be equally likely. As a result, we observe high peaks for methylation frequency throughout the genome, even in areas having lower likelihood ratio positive value as shown in **Figure 3B (i)**, owing to lower coverage in multiple areas. Within the *RDN* locus, we observed few reads showing high positive value of the ratio, but rest of the reads indicated a negative value of the ratio as shown in **Figure 4B (i** and **ii)**. The summary script considered these multiple values as reads and not repeats and thus gave an average value for the locus showing significantly low methylation frequency as shown in **Figure 4B (iii)**.

Another computational tool DeepSignal yielded output in terms of probability of being methylated and probability of being unmethylated at a position in a particular read. If the value of probability of being methylated was higher than 0.5, the position in the read was given a label of methylated and the summary frequency calculation script considered all the values having methylated label to be equally probable. On applying a stringent threshold for probability (>0.9), we observed 5mC predictions only in few regions spread throughout the genome as shown in **Figure 3C (ii)**, interestingly, *RDN* locus being one of these regions. Within the *RDN* locus, we observed few reads showing high 5mC methylation probability, but rest of the reads indicated high probability of unmethylated cytosine as shown in **Figure 4C (i** and **ii)**. The summary script considered these multiple values as reads and not repeats and thus gave an average value for the locus showing significantly low methylation frequency as shown in **Figure 4C (iii)**. This result is in agreement with Nanopolish results and the previous report predicting very few 5mC methylation in CpG motifs of *S. cerevisiae* genome ^[11]^. For the validation of the dataset, we used the raw fast5 data from nanopore sequencing of yeast available on the DeepSignal GitHub repository containing ~4000 R9.4 1D reads basecalled by Albacore (dataset 2), as mentioned before. We observed that the variation with respect to 5mC peak for dampened fraction (Tombo), methylation likelihood (Nanopolish) and probabililty (DeepSignal) was consistent with dataset 1 that is depicted in **Figure 5**. The results for validation experiments are available in the supplementary information.

**Figure 5.**
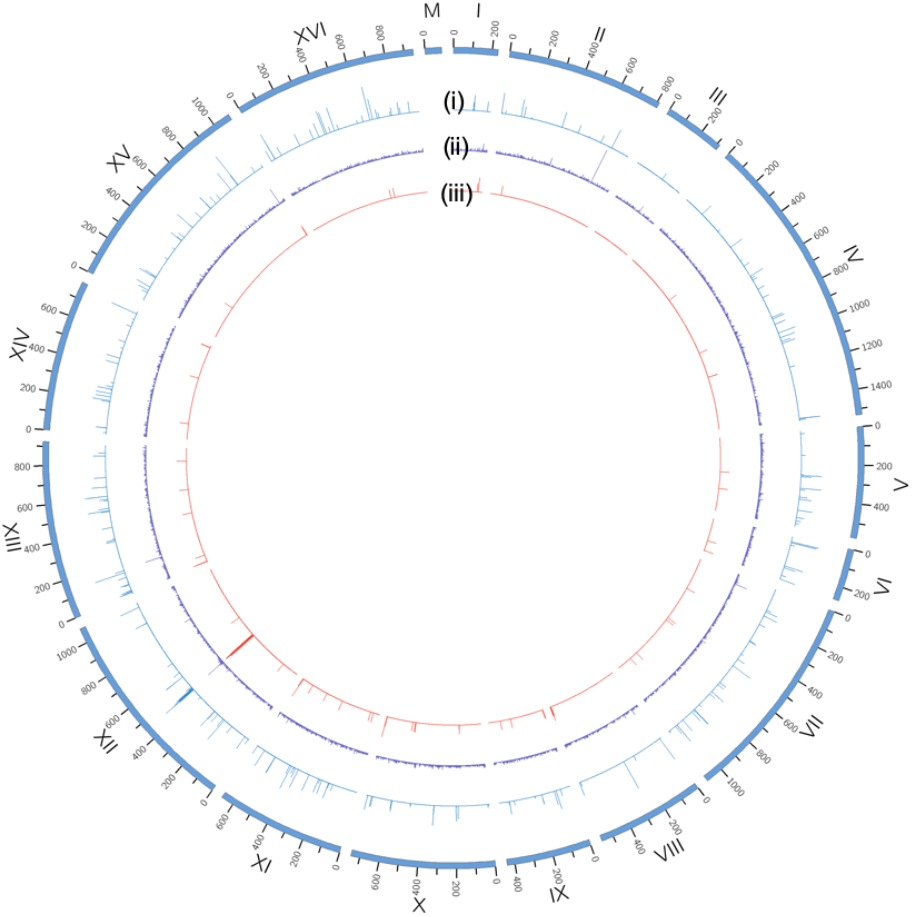
Comparison of 5mC methylation predictions in *S. cerevisiae* genome. Circos plot for comparison of 5mC predictions in *S. cerevisiae* genome showing all chromosomes in a circular map with ticks at every 100kbp and histogram tracks for output data from each computational tool (i) Dampened fraction predicted by Tombo (threshold as >0.5). (ii) Log of Methylation likelihood ratio predicted

The output derived from Nanopolish and DeepSignal for 5mC provided predictions for methylation of CpG motif for every position in every read. Based on a threshold set for each tool, they were called to be methylated or unmethylated and the frequency of methylation for called sites along with coverage was provided for each position whereas Tombo for 5mC provided dampened fraction directly that gave methylation prediction per position after considering the repeated reads. For quantitative analysis **(Table 1),** we calculated a ratio that gave percentage of the motifs predicted to be methylated out of all the motifs present in the sequence according to each computational tool. We performed this analysis for the whole genome as well as for chromosome XII. Nanopolish tool conducted the predictions by considering groups of motifs. As a result, the total number of lines did not correlate to total number of CpG motifs present. Thus, we split the groups into individual CpG motifs and the methylation prediction for the whole group was considered to be true for all the CpG motifs in that group. The values for total number of motif groups, the number of motif groups predicted to be methylated and all the number of values in initial predictions are provided in supplementary Table 1.

For the whole genome in the case of 5mC, we observed that DeepSignal read the maximum number of CpG motifs and gave a low percentage of methylation prediction of 0.94% when the threshold for probability of methylation was set to 0.8, whereas Nanopolish read comparatively lesser number of CpG motifs and yet gave a higher percentage of methylation prediction of 2.8% for both the defined thresholds (1 and 2) (**Table 1**). We observed that Tombo read the least number of motifs and provided the least percentage of methylation prediction of 0.07%. The overall comparison between the tools was consistent for chromosome XII but we observed that Nanopolish and DeepSignal showed an increase in percentage of methylation prediction by approximately 2 folds whereas Tombo showed increase in percentage of methylation prediction by over 8 folds. Our analysis revealed that 5mC predictions with Tombo dampened fraction, Nanopolish methylation likelihood and DeepSignal methylation probability are in agreement with the experimental result with few sites spread throughout the whole yeast genome ^[11, 14]^. The methylation frequency calculation scripts from Nanopolish and DeepSignal overestimated the prediction at regions with low coverage by considering all positive predictions to be equally likely or equally probable and thus require stringent thresholds. At the *RDN* locus, Tombo predicts high methylation fraction whereas Nanopolish and DeepSignal predict high 5mC methylation probability in a smaller fraction of reads and low methylation probability in a larger fraction of reads resulting in overall low methylation frequency. Nanopolish and DeepSignal provide a clearer picture for the repetitive locus by providing predictions of cytosine being methylated as well as unmethylated. At the repetitive *RDN* locus, Tombo shows high methylation fraction whereas Nanopolish and DeepSignal find high probability but low frequency. Since the algorithms do not differentiate between multiple reads and multiple repeats, they tend to give an average value for all repeats which might not be the case if all the repeats are not in the same epigenetic state.

In conclusion, we demonstrated that Tombo and mCaller software tools can indeed be used to predict DNA 6mA modifications. For 5mC, Tombo dampened fraction, Nanopolish methylation likelihood and DeepSignal methylation probability yielded similar results and comparable. This study confirms that 6mA could likely be indeed present in the yeast genome and it is particularly enriched in the *RDN* locus, whereas, 5mC level is rather low throughout the genome including the *RDN* locus.

## Acknowledgement

The work was funded by the Science and Engineering Research Board (SERB), Government of India, (Extra-mural research grant EMR/2016/005135). The fellowship support of ARM and SSP from Ministry of Education, Government of India, is acknowledged.

## Authors’ contributions

RA conceived the study; SSP and ARM performed the experiments and analysis; RA, SSP and ARM wrote the manuscript.

## GitHub Links

1. https://nanoporetech.github.io/tombo/ (for Tombo)
2. https://github.com/al-mcintyre/mCaller (for mCaller)
3. https://github.com/jts/nanopolish (for Nanopolish)
4. https://github.com/bioinfomaticsCSU/deepsignal (for DeepSignal)

