## Supplementary information for "A comparative analysis of computational tools for the prediction of epigenetic DNA methylation from long-read sequencing data"

#### Commands

#### Pipeline for predicting DNA 6mA/5mC modification using Tombo tool

### Installing Tombo on Linux OS via the bioconda environment

```
conda install -c bioconda ont-tombo
```

### Tombo re-squiggling step

```
tombo resquiggle <fast5 files directory> <reference.fasta>
```

### Tombo detect 6mA/5mC modification step

```
tombo detect_modifications alternative_model \
--fast5-basedirs <fast5 files directory>  \
--statistics-file-basename file_name \
--alternate-bases dam
```

```
tombo detect_modifications alternative_model \
--fast5-basedirs <fast5 files directory>\
--statistics-file-basename file_name \
 --alternate-bases CpG
```

### Tombo generating text output files

```
tombo text_output browser_files \
--fast5-basedirs <fast5 files directory> \
--statistics-filename file_name.dam.tombo.stats \
--file-types coverage dampened_fraction \
--browser-file-basename file_name
```

```
tombo text_output browser_files \
--fast5-basedirs <fast5 files directory> \
--statistics-filenamefile_name.CpG.tombo.stats \
--file-types coverage dampened_fraction \
--browser-file-basename file_name
```

### Obtaining a plot with the most significant modified base positions

```
tombo plot most_significant --fast5-basedirs <fast5 files directory> --plot-alternate-model 6mA --statistics-filename file_name.6mA.tombo.stats
```

```
tombo plot most_significant --fast5-basedirs <fast5 files directory> --plot-alternate-model 5mC --statistics-filename file_name.5mC.tombo.stats
```

**Pipeline for predicting DNA 6mA modification using mCaller tool**

### Installing the list of requirements/dependencies for mCaller on Linux OS

```
conda install bwa -y
conda install scikit-learn -y
conda install h5py -y
conda install biopython -y
conda install matplotlib -y
conda install seaborn -y
conda install numpy -y
conda install pysam -y
conda install scipy -y
conda install pandas -y
conda install samtools -y
```

### Downloading and installing mCaller

```
git clone https://github.com/al-mcintyre/mCaller.git
```

### Extracting the template strand reads and saving to .fastq file format

```
nanopolish extract -q -t template <fast5 files directory> -o <filename>.fastq
```

### Indexing the reads

```
nanopolish index -d <fast5 files directory> <filename>.fastq
```

### Aligning the .fastq reads with the reference genome using bwa mem

```
bwa index <reference>.fasta
bwa mem -x ont2d -t <num_threads> <reference>.fasta <filename>.fastq | samtools view -Sb - | samtools sort -T /tmp/<filename>.sorted -o <filename>.sorted.bam
samtools index <filename>.sorted.bam
```

### Eventaligning step

```
nanopolish eventalign -t <num_threads> --scale-events -n -r <filename>.fastq -b <filename>.sorted.bam -g <reference>.fasta > <filename>.eventalign.tsv
```

### mCaller predict 6mA modification step

```
mCaller/mCaller.py -m GATC -r <reference>.fasta -d r95_twobase_model_NN_6_m6A.pkl -e
<filename>.eventalign.tsv -f <filename>.fastq -b A
```

### Generating a summary methylation text .bed file

```
mCaller/make_bed.py -f <filename>.eventalign.diffs.6 -d <min_read_depth> -t <mod_threshold>
```

**Pipeline for predicting DNA 5mC modification using Nanopolish**

### Installing Nanopolish and other requirements on Linux OS via apt install and the bioconda environment

```
sudo apt install nanopolish
conda install minimap2 -y
sudo apt install samtools
```

### Extracting base-calls from fast5 reads and saving into .fastq file

```
nanopolish extract <fast5 files directory> -o output.fastq
```

### Indexing the .fastq file

```
nanopolish index -d <fast5 files directory> output.fastq
```

### Aligning the .fastq file to a reference genome file to create .bam file and indexing it

```
minimap2 -a -x map-ont <reference.fasta> \
samtools sort -T tmp -o output.sorted.bam \
samtools index output.sorted.bam
```

### Calling methylation using the .bam file as input

```
nanopolish call-methylation -t 8 -r output.fastq -b output.sorted.bam -g <reference.fasta> >
methylation_calls.tsv
```

### Generating a methylation summary file in .tsv format

```
python scripts/calculate_methylation_frequency.py methylation_calls.tsv > methylation_frequency.tsv
```

**Pipeline for predicting DNA 5mC modification using DeepSignal**

### Setting up Google Colaboratory for the study

```
! wget https://repo.anaconda.com/miniconda/Miniconda3-py37_4.8.2-Linux-x86_64.sh
! chmod +x Miniconda3-py37_4.8.2-Linux-x86_64.sh
! bash ./Miniconda3-py37_4.8.2-Linux-x86_64.sh -b -f -p /usr/local
import sys
sys.path.append('/usr/local/lib/python3.7/site-packages/')
!conda config --add channels defaults
!conda config --add channels bioconda
!conda config --add channels conda-forge
```

### Installing DeepSignal and other requirements

```
! conda install python=3.6.0 -y
! conda install -c bioconda ont-tombo -y
pip install deepsignal
pip install 'tensorflow==1.12'
```

### Tombo re-squiggling step

```
! tombo resquiggle <fast5 files directory> <reference.fasta>
```

### Extracting signal features around defined motif and saving it as .tsv file

```
! deepsignal extract --fast5_dir <fast5 files directory>
--reference_path <reference.fasta>
--write_path file_name.CpG.signal_features.17bases.rawsignals_360.tsv
```

### Calling modifications using the motif signal features file as input

```
! deepsignal call_mods --input_path /content/file_name.CpG.signal_features.17bases.rawsignals_360.tsv
--model_path
/content/model.CpG.R9.4_1D.human_hx1.bn17.sn360.v0.1.7+/bn_17.sn_360.epoch_9.ckpt
--result_file file_name.CpG.call_mods.tsv
--reference_path  <reference.fasta> --is_gpu no
```

### Generating a methylation summary file in .tsv format

```
! git clone https://github.com/bioinfomaticsCSU/deepsignal.git
! python /content/deepsignal/scripts/call_modification_frequency.py
--input_path /content/file_name.CpG.call_mods.tsv
--result_file file_name.CpG.call_mods.frequency.tsv
--prob_cf 0
```

**Supplementary Table**

| | | Whole Genome | | | | | | |
|---|---|---|---|---|---|---|---|---|
| | | Total number of sites analyzed in all reads | Total readings that are methylated in all reads | Total motif groups that are analyzed | Total motif groups that are methylated | Total CpG or GATC motifs present in the genome sequence | Total CpG or GATC motifs that are methylated in the genome sequence | Ratio of methylated motif to unmethylated motifs (%) |
| **NANOPOLISH** 5mC (CpG) | Log_likelyhood_ratio threshold set to 2 | 662519 | 13892 | 188037 | 6248 | 256946 | 7282 | 2.8341 |
| | Log_likelyhood_ratio threshold set to 3 | 662519 | 7331 | 188037 | 6214 | 256946 | 7244 | 2.8193 |
| **DEEPSIGNAL** 5mC (CpG) | Probability threshold set to 0.8 | 683541 | 3912 | - | - | 387912 | 3674 | 0.9471 |
| **MCALLER** 6mA (GATC) | Probability threshold set to 0.8 | 46392 | 1048 | - | - | 31053 | 936 | 3.0142 |
| | | **Chromosome XII** | | | | | | |
| **NANOPOLISH** 5mC (CpG) | Log_likelyhood_ratio threshold set to 2 | 160876 | 3054 | 16638 | 1162 | 23042 | 1440 | 6.2495 |
| | Log_likelyhood_ratio threshold set to 3 | 160876 | 1568 | 16638 | 924 | 23042 | 1141 | 4.9518 |
| **DEEPSIGNAL** 5mC (CpG) | Probability threshold set to 0.8 | 142276 | 787 | - | - | 34617 | 587 | 1.6957 |
| **MCALLER** 6mA (GATC) | Probability threshold set to 0.8 | 7516 | 184 | - | - | 2677 | 104 | 3.8849 |

**Supplementary Table 1. Detailed quantitative analysis of methylation predictions using computational tools.** The table gives a detail summary of the total number of motifs that were read to be present and how many of those motifs were predicted to be methylated according to each computational tool along with a ratio of both the quantities in percentage. We performed this analysis for the whole genome as well as for chromosome XII. As additional information, here we show the total derived number of sites analyzed in all reads and how many of those sites were methylated as predicted by DeepSignal, Nanopolish and mCaller. Nanopolish tool conducted the predictions by considering groups of motifs. We derived the total number of motif groups that are present and how many of those groups are predicted to be methylated.

**Supplementary Figure**

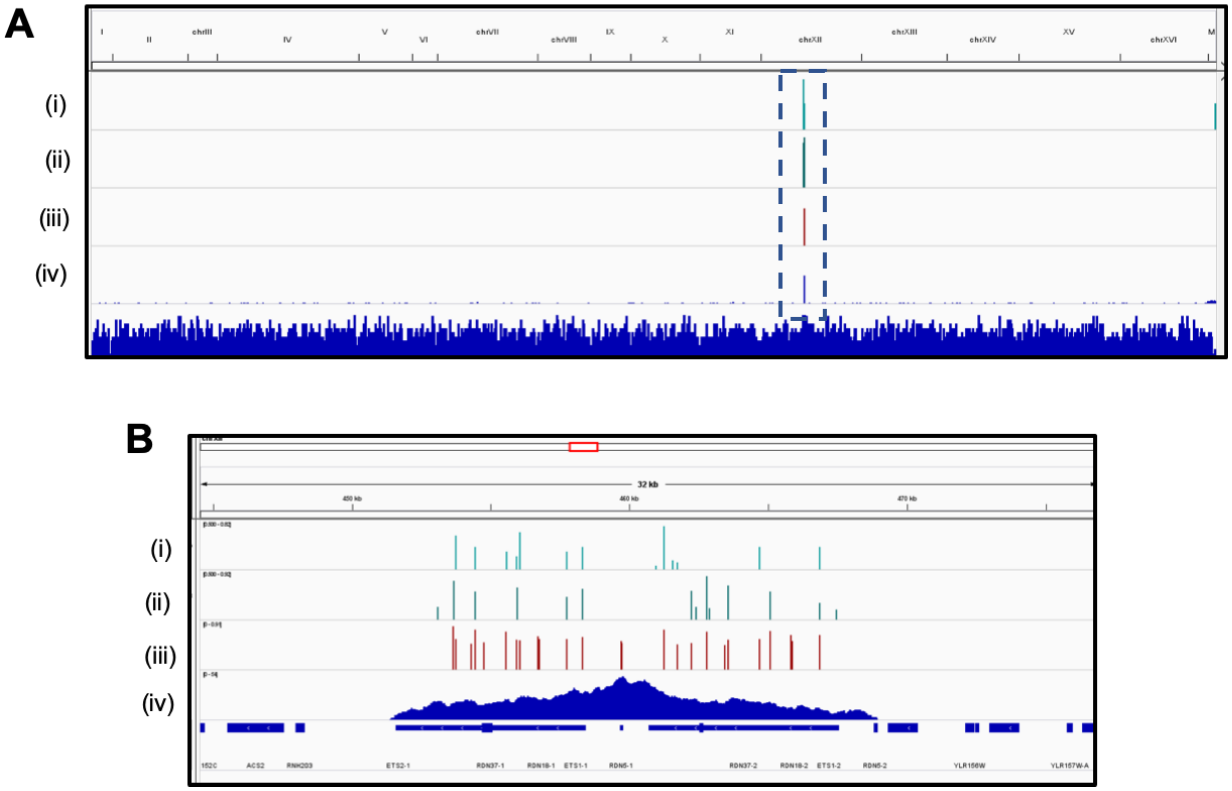


**Supplementary Figure 1. Validating the prediction of 6mA modification in *S. cerevisiae* using dataset 2. A**. Prediction for the whole genome. On the top, all the chromosome details are displayed and on the bottom, we have the collapsed gene view of SacCer3 for comparison of the whole genome. (i) Track showing high DNA 6mA prediction by Tombo, for forward strand, (threshold >0.5), (ii) Track showing high DNA 6mA prediction by Tombo, for reverse strand (threshold >0.5), (iii) Track showing high DNA 6mA prediction by mCaller (threshold >0.5). (iv) Track showing high coverage for 6mA prediction by Tombo. The box (- - -) indicates region of high dampened fraction on chromosome XII. **B.** Prediction for the RDN locus. In IGV, the red rectangular box on top indicates the region of chromosome XII that is zoomed in with its size written below it and the expanded form of SacCer3 gene is displayed below. (i) Track showing high 6mA prediction for forward strand, by Tombo. (threshold >0.5) (ii) Track showing high 6mA prediction for reverse strand, by Tombo. (threshold >0.5) (iii) Track showing high 6mA prediction for mCaller. (threshold >0.5) (iv) Track showing high coverage for 6mA prediction by Tombo.

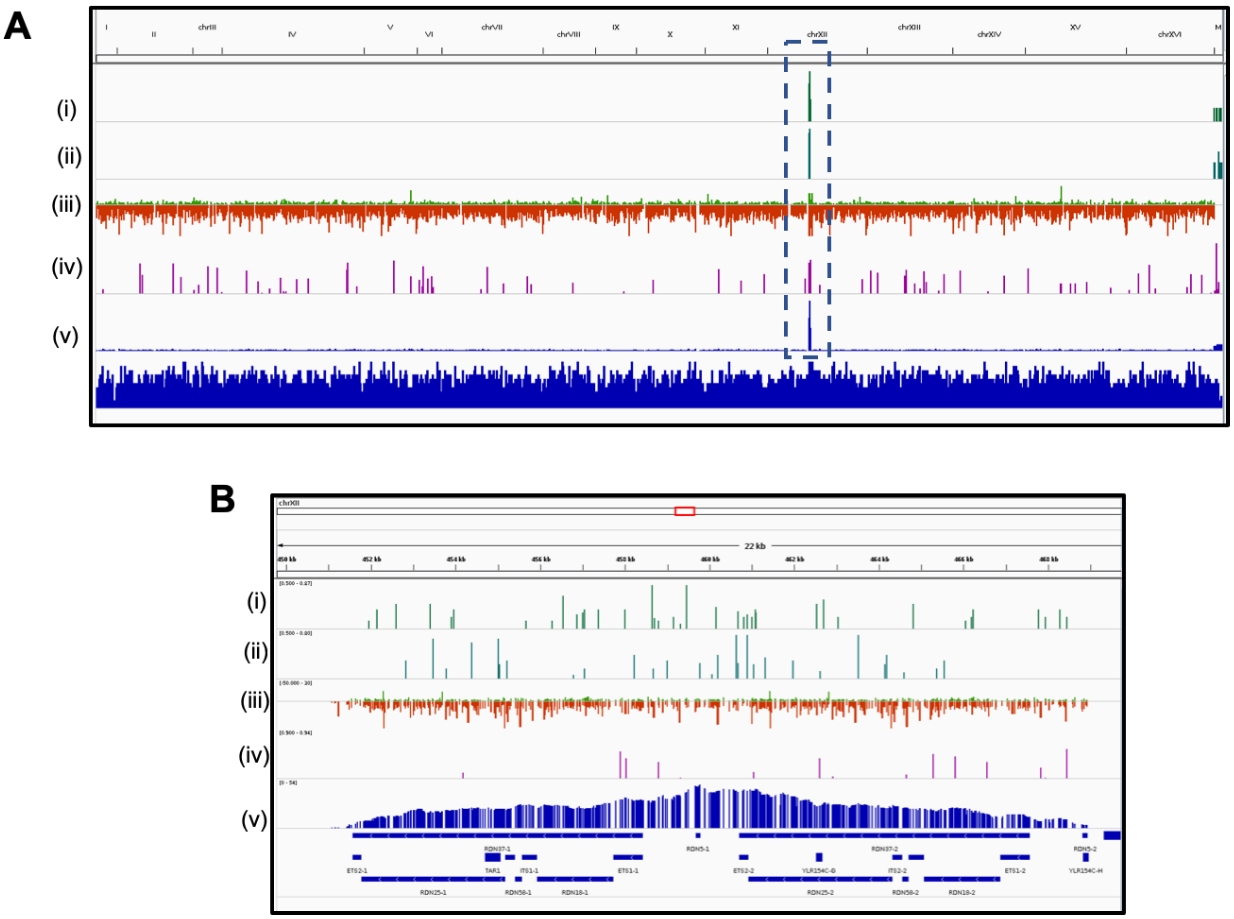


**Supplementary Figure 2. Validating the prediction of 5mC modification in *S. cerevisiae* using dataset 2. A**. Prediction for the whole genome. At the top, all the chromosome details are displayed and at the bottom, we have the collapsed gene view of SacCer3 for comparison of the whole genome. (i) Track showing DNA 5mC prediction by Tombo, for forward strand (threshold >0.5), (ii) Track showing DNA 5mC prediction by Tombo, for reverse strand (threshold >0.5), (iii)Track showing log likelihood ratio of 5mC modification predicted by Nanopolish (-50 to 30) with positive values in green and negative values in red. (iii) Track showing 5mC Methylation probability predicted by DeepSignal (threshold >0.9), (v) Track showing coverage as calculated using DeepSignal. The box (- - -) indicates region of high dampened fraction on chromosome XII**. B.** Prediction for the RDN locus. In IGV, the red rectangular box on top indicates the region of chromosome XII that is zoomed in with its size written below it and the expanded form of SacCer3 gene is displayed below. . (i) Track showing DNA 5mC prediction by Tombo, for forward strand (threshold >0.5), (ii) Track showing DNA 5mC prediction by Tombo, for reverse strand (threshold >0.5), (iii)Track showing log likelihood ratio of 5mC modification predicted by Nanopolish (-50 to 30) with positive values in green and negative values in red, (iii) Track showing 5mC Methylation probability predicted by DeepSignal (threshold >0.9), (v) Track showing coverage as calculated using DeepSignal.
